# CaLMPhosKAN: Prediction of General Phosphorylation Sites in Proteins via Fusion of Codon-Aware Embeddings with Amino Acid-Aware Embeddings and Wavelet-based Kolmogorov–Arnold Network

**DOI:** 10.1101/2024.07.30.605530

**Authors:** Pawel Pratyush, Callen Carrier, Suresh Pokharel, Hamid D. Ismail, Meenal Chaudhari, Dukka B. KC

## Abstract

The mapping from codon to amino acid is surjective due to the high degeneracy of the codon alphabet, suggesting that codon space might harbor higher information content. Embeddings from the codon language model have recently demonstrated success in various downstream tasks. However, predictive models for phosphorylation sites, arguably the most studied Post-Translational Modification (PTM), and PTM sites in general, have predominantly relied on amino acid-level representations. This work introduces a novel approach for prediction of phosphorylation sites by incorporating codon-level information through embeddings from a recently developed codon language model trained exclusively on protein-coding DNA sequences. Protein sequences are first meticulously mapped to reliable coding sequences and encoded using this encoder to generate codon-aware embeddings. These embeddings are then integrated with amino acid-aware embeddings obtained from a protein language model through an early fusion strategy. Subsequently, a window-level representation of the site of interest is formed from the fused embeddings within a defined window frame. A ConvBiGRU network extracts features capturing spatiotemporal correlations between proximal residues within the window, followed by a Kolmogorov-Arnold Network (KAN) based on the Derivative of Gaussian (DoG) wavelet transform function to produce the prediction inference for the site. We dub the overall model integrating these elements as CaLMPhosKAN. On independent testing with Serine-Threonine (combined) and Tyrosine test sets, CaLMPhosKAN outperforms existing approaches. Furthermore, we demonstrate the model’s effectiveness in predicting sites within intrinsically disordered regions of proteins. Overall, CaLMPhosKAN emerges as a robust predictor of general phosphosites in proteins. CaLMPhosKAN will be released publicly soon.

## 1. Introduction

A post-translational modification (PTM) is a chemical alteration in proteins that occurs after translation from mRNA, contributing significantly to molecular diversity and functional dynamics within cells ^1^. These modifications are crucial in regulating essential cellular processes and altering protein functions. Among PTMs, phosphorylation is particularly well-studied ^2^ and involves the covalent addition of a phosphate group to the N-linked amino acid chain or side chain, typically mediated by kinase enzymes ^3^. This modification is most commonly observed on the residues of Serine (S), Threonine (T), and Tyrosine (Y), though it also occurs on Aspartic acid ^4^, arginine ^5^, cysteine ^6^, and histidine^7,8^. Phosphorylation serves as a fundamental regulatory mechanism in various biological processes, including muscle contraction, cell growth, neural activity, and signal transduction ^9^. The dysregulation of phosphorylation, potentially induced by natural toxins or pathogens, can lead to severe diseases such as cancer, Alzheimer’s, and heart disease ^10^. Consequently, the identification and understanding of phosphorylation sites are critical for developing new therapeutic strategies and insights into drug design ^11–13^.

Phosphorylated proteins can be experimentally identified through methods such as P-labeling and mass spectrometry ^14,15^, which enable the labeling of each specific residue within a peptide as either phosphorylated or non-phosphorylated. However, these techniques are often time-consuming and costly. Additionally, they cannot keep pace with the rapid availability of sequencing data, thereby widening the gap between the function annotation and the burgeoning number of sequences. In contrast, computational methods leveraging machine learning and deep learning offer a more efficient and effective tool for various bioinformatics tasks ^16^ including phosphorylation site prediction. These methods typically involve converting protein peptide or FASTA sequences into feature vectors, which are then used to predict the phosphorylation status of specific residues within the sequence. Traditionally, computational models for general phosphorylation site prediction are categorized into two types: an *S+T* model and a *Y* model, reflecting the differences in the residues where the phosphorylation occurs. However, recent developments, such as a model for prokaryotic phosphorylation ^17^ based on capsule networks, have highlighted the need for three distinct models, one each for Serine (S), Threonine (T), and Tyrosine (Y). This differentiation highlights the importance of tailoring computational methods to the specific characteristics and patterns within the sequences.

In recent years, there has been a noticeable shift from traditional machine learning-based models to more advanced deep learning-based models for predicting PTMs in proteins. Machine learning methods such as RFPhos ^18^, NetPhosK ^19^, KinasePhos ^20^, and GPS ^21^ relied on manually extracted features, including physiochemical properties, disordered regions, and other biological information to model phosphorylation. While these models provided acceptable prediction capabilities, they were generally outperformed by the more sophisticated deep-learning approaches ^22–25^. Deep learning methods in this field typically utilize learned embeddings or leverage pre-trained language models to enhance prediction accuracy. One of the pioneering deep learning models, Musite ^26^, introduced the use of attention-based mechanisms to improve the prediction of phosphorylation PTMs. Building on this, the same research group developed CapsNet^17^, which employed a capsule network architecture. This model not only marginally improved upon its predecessor but also enhanced the interpretability of the predictions through the use of capsules. Another important advancement came with DeepPSP ^27^, which expanded upon earlier models by incorporating global contextual information around potential phosphorylation sites, rather than focusing solely on local sequence information. DeepPSP integrated this global context through the use of Squeeze-and-Excitation Networks (SENet), moving towards demystifying the often black-box nature of deep learning models. Most recently, LMPhosSite ^28^ has utilized residue-level embeddings from pre-trained protein language models (pLMs), marking a further evolution in the computational prediction of phosphorylation sites. While considerable advancements have been made in developing computational tools for phosphorylation site prediction, there remains ample scope for enhancing their predictive performance. Moreover, all these tools have traditionally relied on representations derived from amino-acid sequences. Recently, Outeiral *et al.* ^29^ introduced CaLM (Codon Adaption Language Model), a protein language model trained on protein-coding DNA sequences. They observed that the codon pLM CaLM outperforms amino acid-based state-of-the-art pLMs, such as ESM ^30^, ProtT5 ^31^, and ProtBERT ^31^, in various tasks including melting point prediction, solubility prediction, subcellular localization classification, and function prediction ^32–34^.

In this study, we explore the potential of codon-aware embeddings for general phosphorylation site prediction. We expand the contemporary approaches that use amino acid-aware protein language models by integrating embeddings from CaLM, aiming to capture potentially higher information content inherent in the codon space. Specifically, we combine these codon-aware embeddings from CaLM with amino acid-aware embeddings from the ProtTrans encoder to create a merged representation of the full sequence. Subsequently, we extract features that capture the spatiotemporal correlations of residues at the window level, which are learned by a wavelet-based Kolmogorov–Arnold Network (KAN) model. Our model, dubbed CaLMPhosKAN, outperforms existing phosphorylation site predictors, demonstrating that integrating codon-based embeddings can enhance general phosphosite prediction. To the best of our knowledge, this is the first work to utilize codon-aware embeddings in PTM prediction problems.

## 2. Materials and Methods

### 2.1 Dataset Construction

In this study, we use the dataset of Li *et al.*, comprising experimentally identified phosphorylation sites curated from multiple databases ^27^. To ensure diversity, the protein sequences underwent homology reduction using CD-HIT ^35^ with a similarity cut-off of 0.5. The experimentally annotated serine (S), threonine (T), and tyrosine (Y) residues were designated as phosphorylated sites (P-sites). In contrast, the remaining non-annotated S, T, and Y residues within the same sequences were considered non-phosphorylated (NP-sites). Following this, the dataset was randomly divided into training and test sets in a 9:1 ratio based on protein IDs instead of sites, ensuring no overlap of sequences between the sets. Such a division is crucial because it prevents homology between the sets and contextual information leakage from train to test. This is particularly important since our work leverages protein language models that operate on the full sequence context. We excluded a few proteins from our dataset that had been updated in the UniProt database (timestamp: May 2024) and no longer matched their original annotations.

The subsequent step involved mapping the protein sequences to their corresponding coding DNA sequences, a process characterized by non-injective mapping (or surjective mapping if considered in the reverse direction, see Figure 1). We initiated this by generating protein-coding sequences through the mapping of UniProt identifiers to NCBI nucleotide reference sequences (RefSeq). These RefSeqs are curated to be non-redundant and reliable representations of nucleotide sequences. We used the accession numbers of the identified RefSeqs to retrieve their corresponding candidate coding sequences from the NCBI Nucleotide database ^36^ (timestamp: May 2024), specifically in GenBank format. This format provides comprehensive details, including the coding sequences and the translated protein product. To ensure the accuracy of the coding sequences as the correct coding sequence for each mapped UniProt protein, we performed a pairwise global alignment using the Needleman–Wunsch (NW) algorithm ^37^ between the translated protein sequence from the GenBank format and the original UniProt sequence. The identity percent score for each alignment was computed using the NW algorithm. For inclusion in our dataset, we selected only those coding sequences where the alignment demonstrated 100% identity with the UniProt protein sequence. Proteins that exhibited less than 100% alignment identity were excluded. Additionally, proteins for which corresponding RefSeqs were unavailable were discarded from the dataset.

**Figure 1.**
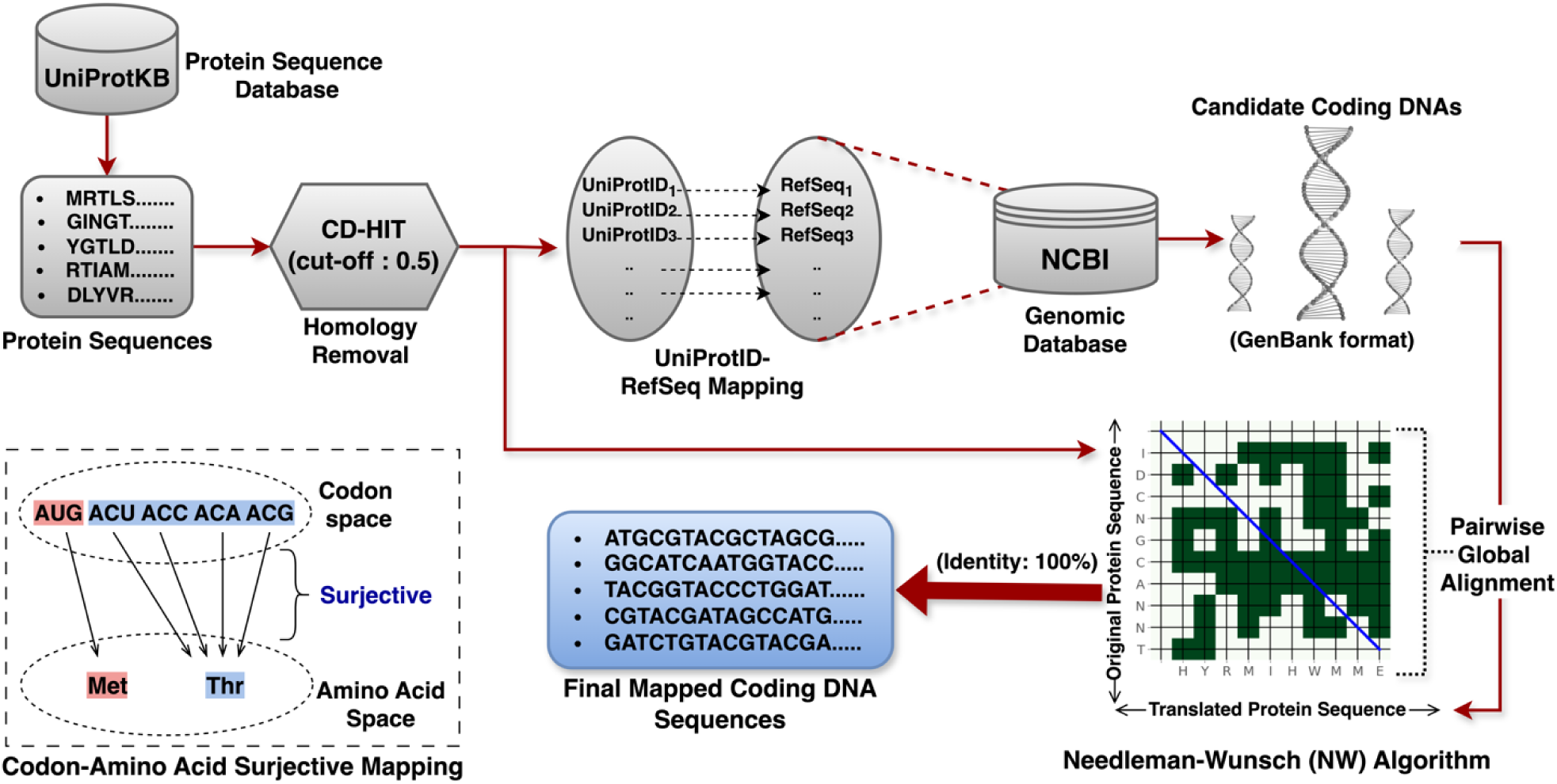
A high-level overview of the dataset preparation and the mapping procedure for obtaining protein-coding DNAs. The lower left corner illustrates the nature of codon alphabets mapping to amino acid alphabets, showing a surjective mapping. For instance, the amino acid Methionine (Met, *M*) is mapped by a single codon ‘AUG’, while Threonine (Thr, *T*) can be mapped by at most four codons: ‘ACU’, ‘ACC’, ‘ACA’, and ‘ACG’.

Finally, the termination codons (stop codons) ‘TAA’ (Ochre), ‘TAG’ (Amber), and ‘TGA’ (Opal/Umber) were removed from each coding sequence, resulting in the coding DNA sequence length being exactly thrice the length of the corresponding protein sequence, aligning with the triplet nature of the genetic code. The high-level overview of the whole procedure is depicted in Figure 1.

Table 1 summarizes the final processed data for *S*, *T,* and *Y* residues, including the number of phosphorylated sites (P-sites) and non-phosphorylated sites (NP-sites), the count of coding DNAs/proteins, the mean length with one standard deviation, and the ratio of NP to P. Given the structural and functional similarities between serine (S) and threonine (T) residues, we combined their datasets. Both serine (S) and threonine (T) possess a hydroxyl group (-OH) on their side chains, making them similarly accessible and reactive to the same kinases for phosphorylation ^38^. In contrast, tyrosine (Y) residues are subject to phosphorylation through distinct enzymatic processes due to the presence of *π*-electrons and conjugated electron systems in tyrosine’s phenol ring creates unique interactions with tyrosine kinases, differing fundamentally from the way serine and threonine interact with their respective kinases ^39^. Due to these differences in enzymatic mechanisms, we maintained a separate dataset for *Y* sites, distinct from the combined *S/T* (or *S+T*) dataset. The training sets in both datasets (*S+T* and *Y*) were balanced using random undersampling to avoid biased modeling while the distribution of targets in the independent test was left unaltered for evaluating generalization loss. Reflecting on many works in phosphorylation prediction ^17,19–21,27,28^, we developed two distinct models: one for the combined *S+T* residues and another specifically for *Y* residues.

**Table 1.** Description of the training set (before balancing) and independent testing set respective to each target residues - Serine (S), Threonine (T), and Tyrosine (Y) residues.

| Set | Target Residue | #coding<br>DNA/protein | Coding<br>DNA length | P-sites<br>(+ve) | NP-sites<br>(-ve) | Ratio<br>(NP:P) |
| --- | --- | --- | --- | --- | --- | --- |
| Training | Serine (S) | 11,363 | 1847±1540 | 110,752 | 479,531 | 4.33:1 |
|  | Threonine (T) | 11,355 | 1848±1540 | 43,468 | 320,798 | 7.38:1 |
|  | Tyrosine (Y) | 8178 | 2063±1673 | 27,077 | 123,918 | 4.57:1 |
|  | **Total** |  |  | **181,297** | **924,247** | **5.09:1** |
| Ind. Testing | Serine (S) | 1248 | 1806±1396 | 12,077 | 50,492 | 4.18:1 |
|  | Threonine (T) | 1246 | 1808±1369 | 4887 | 33,565 | 6.87:1 |
|  | Tyrosine (Y) | 902 | 2062±1711 | 3054 | 13,347 | 4.37:1 |
|  | **Total** |  |  | **20,018** | **97,404** | **4.90:1** |

### 2.2 Protein Embeddings

The highly degenerate nature of codons leads to a surjective mapping from multiple codons to a single amino acid, with most amino acids being encoded by up to six different codons (see Figure 1). This indicates that a sequence represented at the codon level might contain as much or more information than the same sequence at the amino acid level. To harness this potential, input sequences are represented using both informational modes: codon-level and amino-acid level. This approach generates a richer sequence representation by combining embeddings from both domains, each produced by protein language models (pLMs) pre-trained on their respective modal representations. Codon-level embeddings, derived from a pLM trained on coding DNA sequences, are designed to capture translational nuances and gene expression influences that are specific to certain codon arrangements ^40^. Conversely, amino acid-level embeddings, obtained from a pLM trained on protein sequences, primarily reflect the functional potentials of proteins based on their amino acid composition ^41^. A detailed explanation of these embeddings is provided below.

#### 2.2.1 Codon-Aware Embeddings

The coding DNA sequences are encoded using a specialized codon-aware protein language model called CaLM (Codon adaptation Language Model) ^29^. Built on the Evolutionary Sequence Modelling (ESM) framework, CaLM utilizes an architecture comprising 12 encoder layers (each with 12 attention heads) and a prediction head, amounting to 86 million parameters in total. This model undergoes pretraining using a masked language modeling denoising objective on a dataset of approximately 9 million non-redundant coding sequences derived from whole-genome sequencing.

Prior to encoding input coding sequences, the sequences are tokenized into integer tokens that map to the 64 codons ‘words’, along with special tokens. The special tokens <CLS> (classification) and <EOS> (end-of-sequence) are added at the start and end of the sequence, respectively. The vectorized tokens are then fed into the encoder, and the last hidden states (also called embeddings) are extracted. For an input coding sequence of length 3*N* (corresponding to a protein sequence of length *N*), an embedding matrix of dimension (*N*+2) x *L* is generated, where *L* is the embedding dimension (=768). It is noteworthy that although the input coding sequence is three times the length of the corresponding protein sequence, the resulting embedding matrix corresponds to the length of the protein sequence. This tensor includes the embeddings for the special tokens <CLS> and <EOS> which are discarded, resulting in an *N* x 768-dimensional matrix representing the input sequence.

#### 2.2.2 Amino Acid-Aware Embeddings

Amino acid-aware embeddings are derived from a protein language model trained on a large corpus of protein sequences. In this work, we utilize a ProtTrans family model called ProtT5 ^31^, a prominent pLM established for its high performance in various protein downstream tasks^42–44,44^, including post-translational modification prediction ^24,25,28,32–34,45,46^. ProtT5 is built on the T5 (Text-to-Text Transfer Transformer)^47^ architecture and has been trained using MLM denoising objective on the UniRef50 ^48^ (UniProt Reference Clusters, encompassing 45 million protein sequences) database. The model comprises a 24-layer encoder-decoder architecture (each with 32 attention heads) and contains approximately 2.8 billion learnable parameters. For this work, we employed the pre-trained encoder component of ProtT5 to extract embeddings from the input protein sequences.

For a given protein sequence of length *N*, each amino acid is converted into integer tokens that map to a vocabulary of 21 canonical amino acids plus a special <EOS> token. Non-canonical amino acids such as ‘U’ (Selenocysteine), ‘Z’ (Pyrrolysine), ‘O’ (Hydroxyproline), and ‘B’ (Beta-amino acids) are mapped to a pseudo-amino acid represented as ‘X’. The tokenized sequence, which now consists of these integer tokens, is then processed through the encoder’s attention stack of the ProtT5 model. From this, we extract the last hidden state, which has a dimension of 1024. As a result, with an input protein sequence of length *N*, ProtT5 produces amino-acid context-aware embeddings with dimensions (*N*+1) × *L*, where *L* is 1024. The embedding vector corresponding to the <EOS> token is discarded, and the remaining *N*×1024 dimensional tensor is used for downstream learning.

### 2.3. Proposed Architecture

The architecture of CaLMPhosKAN is structured into three interconnected modules, each designed to handle specific functions within the prediction framework. The complete schematic of CaLMPhosKAN is depicted in Figure 2. The ‘Embedding Extraction and Fusion Module’ generates and integrates codon-level and amino-acid-level contextualized embeddings from protein language models. The ‘Spatiotemporal Feature Extraction Module’ then extracts crucial features that capture spatiotemporal correlations within local windows around the site of interest. Finally, the ‘Wav-KAN Module (or Classification Module)’ employs a wavelet-induced KAN model to classify these features into P sites and NP sites. Note that while the overall architecture remains consistent for *S+T* and *Y* datasets, the model implementation differs only in the Wav-KAN configuration, a detail that will be discussed later. An elaborated explanation of these three modules is presented below.

#### 2.3.1 Embedding Extraction and Fusion Module

This module integrates multiple components: encoders, a mapper, an early fusion mechanism, and window-level embedding extractor. Initially, the full protein sequence of length *N* is processed using the ProtTrans encoder, which transforms the sequence into an *N*×1024-dimensional embedding matrix. Concurrently, the mapper converts this *N*-length full protein sequence into its corresponding 3*N* length coding DNA sequence (details in section 2.1). The translated coding sequence is then embedded using the CaLM encoder, resulting in an *N*×768-dimensional embedding matrix. For the fusion step, the *N*×1024-dimensional matrix from the ProtTrans encoder and the *N*×768-dimensional matrix from the CaLM encoder is concatenated horizontally. This early fusion results in an *N*×1792-dimensional matrix, which integrates amino acid and codon-level information, providing a rich representation of the input sequence.

To represent the site of interest (*S*/*T*/*Y*) within the protein, we define a window frame of size *W* around the tokens corresponding to the sites. This window includes ⎣*W*/2⎦ upstream and downstream flanking residues, with the site of interest located at position ⎣*W*/2⎦ + 1 (one-based indexing). The fused embeddings within this window frame, constituting a *W*×1792-dimensional matrix, will represent the site of interest. Based on cross-validation experiments and computational efficieny, the optimal value of *W* was determined to be 9 (four resides on each side with the 5^th^ residue as the site of interest).

**Figure 2.**
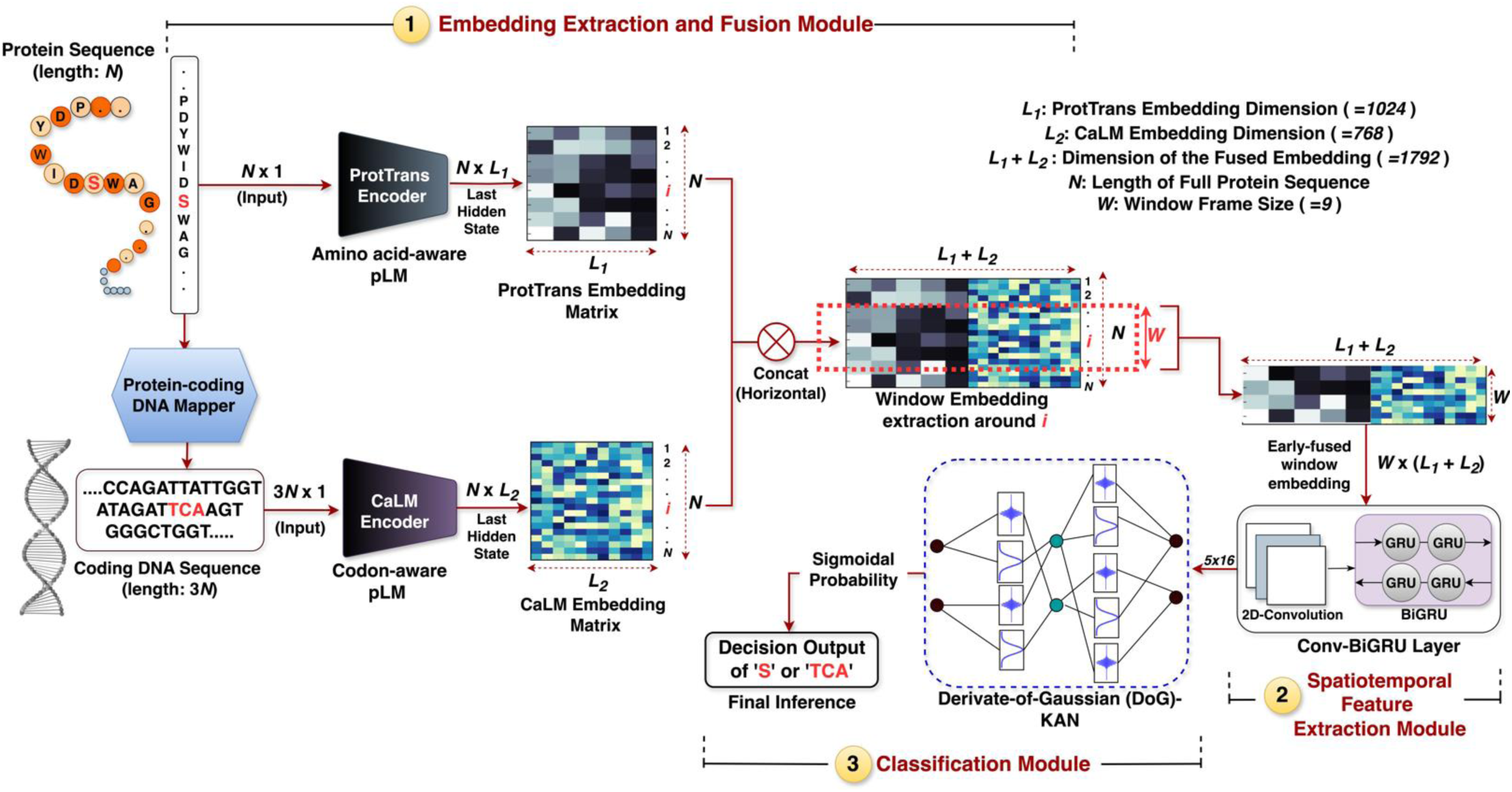
The architecture of CaLMPhosKAN depicting the three modules - (1) Embedding Extraction and Fusion Module, (2) Spatiotemporal Feature Extraction Module, and (3) Wav-KAN Module (or, Classification Module). The site of interrogation in the protein sequence is ‘S’ (at index *i*) or ‘TCA’ (at index 3*i*) in the corresponding coding DNA, and is highlighted in bold red. The input is the full-length protein sequence containing the site, provided to the Embedding Fusion and Extraction Module, and the output is the decision inference of the site (S/TCA) obtained from the Wav-KAN Module.

#### 2.3.2 Spatiotemporal Feature Extraction Module

Kinases, which catalyze the phosphorylation process, often recognize specific sequences or motifs near the phosphorylation site. Additionally, the neighboring residues can induce conformational changes that either expose or hide potential phosphorylation sites, impacting their accessibility ^49^. To capture these intricate interactions, our window-level feature extraction module leverages correlations among neighboring residues within a specified window frame around the site of interest.

To extract spatial correlations, we utilize a 2D-convolutional layer. The input to this layer is a matrix of dimension 1792×9, where 1792 is the dimension of fused embeddings and 9 is the size of the window frame around the site. The layer applies 16 kernels of size 5×5 across this matrix with a single channel, producing a 16×5×1788 dimensional tensor corresponding to the window frame. Following the spatial feature extraction, a Bidirectional Gated Recurrent Unit (BiGRU) layer is employed with 8 units to capture the sequential context within the window frame. The BiGRU layer processes the sequence in both forward and reverse directions, learning dependencies from residues preceding and following the target. The output dimension of the ConvBiGRU layer is 5×16 which serves as an input to the classification module (Wav-KAN) which is described in the next subsection.

#### 2.3.3 Wav-KAN Module

The Wav-KAN module, or classification module, incorporates a Wav-KAN (Kolmogorov Arnold Network) network ^50^ tasked with rendering the final classification inference. This process begins with the input with dimension 5×16 from the ConvBiGRU layer, which is flattened into an 80×1 length vector before being introduced into the Wav-KAN network. Two distinct Wav-KAN models are tailored for the two target residue-specific datasets: one for the *S+T* and another for the *Y* datasets. The model designed for the *S+T* dataset features a KAN with two hidden layers; the first contains 128 nodes and the second contains 32 nodes. In contrast, the model for the *Y* dataset is simpler, utilizing a single hidden layer with 24 nodes. A batch normalization layer precedes each hidden layer in both *S+T* and *Y* models.

Unlike traditional Multi-Layer Perceptrons (MLPs) that use fixed activation functions and linear weights at nodes, the KAN architecture utilizes learnable univariate functions on each edge, which are then summed across the nodes of subsequent layers. Furthermore, Wav-KAN is a variation of KAN ^51^ chosen for its ability to enhance performance and reduce training time by incorporating wavelet transformation functions as approximate activation functions. These wavelets transform input from each node along the model’s edges through a defined and trainable function known as the ‘mother wavelet’. The Wav-KAN supports multiple mother wavelet functions, including both Continuous Wavelet Transform (CWT) and Discrete Wavelet Transform (DWT). For this work, we selected the Derivative of Gaussian (dubbed “DoG”) wavelet function based on its performance in 10-fold cross-validation. The DoG wavelet can be defined by Equation 1.

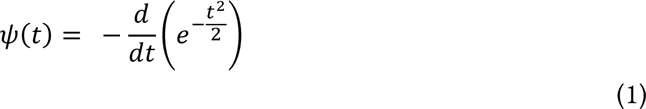

where *ψ(t)* represents the wavelet function dependent on the time variable *t.* This mother wavelet function is further trainable with the Equation 2 given below:

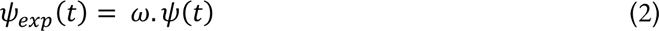

where *w* serves as a learnable coefficient for the mother wavelet function, enabling fine-tuning of the wavelet shape during training.

Following the hidden layers, the output passes through a single neuron equipped with a sigmoid activation function, converting the output into probabilities ranging from 0 to 1. These probability values are then used to make the classification inference, determining whether the site of interrogation (marked as ‘S’ in the protein sequence and ‘TCA’ in the corresponding coding DNA sequence in Figure 2) belongs to a P or NP site. A detailed architectural description of the ConvBiGRU with Wav-KAN integration is provided in Supplementary Section S1. Note that the optimization of the ConvBiGRU and Wav-KAN models were performed using a 10-fold cross-validation.

### 2.4. Training and Evaluation Protocol

The proposed model is trained to minimize binary-cross entropy with logits (BCEwithLogits) loss function using the Adam optimizer. This loss function can be mathematically defined as:

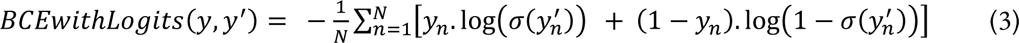

where *N* is the number of observations in the batch, *y_n’_*is the ground truth, and *y_n_*`’` is the logit from the model for the observation *n*. σ(*y_n_’*) is the sigmoid function applied to the logit *y_n_*’ given by:

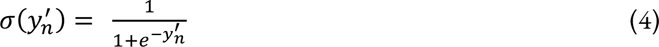

The training is conducted in a mixed precision floating point (leveraging both 16-bit and 32-bit operations) to optimize computational efficiency and memory usage. The loss is dynamically scaled during backpropagation using Pytorch’s GradScaler ^52^ to ensure the gradients are sufficiently large to avoid underflow when using 16-bit precision. The adaptive learning rate of 8e^-4^ is chosen with a decay rate of 0.9 for the first moment 0.999 for the second moment and a batch size of 1024. The optimization of hyperparameters is performed using stratified 10-fold cross-validation on the training set, ensuring that proteins in each training fold are mutually exclusive with those in the corresponding validation set. Early stopping is employed to avoid overfitting, and accuracy/loss curves (see sample curves in Supplementary Section S3) are carefully monitored in each fold. Additionally, a model checkpoint is used to save the best model over the epochs. It is worth noting that deep learning models (including pLMs) are implemented in a PyTorch environment, with training and inference conducted on an NVIDIA A100-SXM4-80GB GPU.

Model evaluation is conducted using five performance metrics consistent with existing works ^27^. These metrics include Matthews Correlation Coefficient (MCC), Precision (PRE), Recall (REC), F1, and Area Under Curve (AUC). Due to the imbalance in our independent test set, we use the weighted version of the F1-score i.e. F1_weighted_ and Area Under the Precision-Recall Curve (AUPR) instead of the Area Under Receiver Operating Characteristic (AUROC) ^53,54^. A detailed description of these metrics is presented in Supplementary Section S2.

Additionally, the statistical significance of our proposed model against other approaches is computed using McNemar’s chi-squared test with a continuity correction ^55^, set at a standard significance level (***α***) of 0.05. The null hypothesis in this case, which states that the two models are statistically similar, is rejected if the *p*-value is below the significance level.

## 3. Results

In this section, we present various empirical studies performed to evaluate CaLMPhosKAN. We first provide the cross-validation analysis conducted on various embeddings and multiple wavelet functions of the KAN model. Following this, we compare our proposed model, CaLMPhosKAN, with existing state-of-the-art methods. Finally, we assess the predictive ability of CaLMPhosKAN on Intrinsically Disordered Regions (IDRs) and non-Intrinsically Disordered Regions (non-IDRs) of proteins.

### 3.1. Evaluation of Embeddings

We aim to assess the contribution of codon-aware embeddings to the final predictive performance of CaLMPhosKAN. To this end, we perform 10-fold cross-validation (see Table 2) independently on CaLM embeddings (codon-aware), ProtTrans embeddings (amino acid-aware), and the fused embeddings (CaLM + ProtTrans). On the *S+T* dataset, CaLM embeddings produced a mean MCC, mean F1_weighted_, and mean AUPR of 0.44±0.01, 0.71±0.01, and 0.80±0.01 respectively, which is worse than ProtTrans embeddings, which produced a mean MCC, mean F1_weighted_, and mean AUPR of 0.46±0.01, 0.72±0.01, and 0.81±0.01 respectively. However, upon combining the two sets of embeddings via early fusion (aka CaLMPhosKAN), we observed an improvement in performance metrics with a mean MCC, mean F1_weighted_, and mean AUPR of 0.48±0.01, 0.74±0.01, and 0.83±0.01. Similarly, on the *Y* dataset, CaLM embeddings alone did not outperform ProtTrans embeddings. Yet, the combination of the two through early fusion resulted in better performance metrics than when either pLM was used independently. This improvement upon integration suggests that codon-aware embeddings contribute complementary information that enhances the overall model performance.

Interestingly, although CaLM did not independently surpass ProtTrans in performance in both datasets, the differences were not substantial. This is particularly notable considering that CaLM operates with roughly 33 times fewer parameters, highlighting its ability to deliver considerable predictive value with substantially reduced model complexity.

To further understand the observed improvement when integrating codon-aware and amino acid-aware embeddings, we analyzed the attention weights from both the CaLM and ProtTrans protein language models (pLMs). Figure 2 illustrates heatmaps generated from the final layer of the encoder stack in both pLMs, averaged across all attention heads for a complete protein sequence (ID: P30047) with 86 tokens excluding special tokens. Note that the heatmaps of individual attention heads (32 per layer in ProtTrans and 12 per layer in CaLM) from the final encoder are also provided. Heatmaps for ProtTrans can be found in Supplementary Section S4, while those for CaLM are presented in Figure 2c. These heatmaps indicate a strong association between neighboring residues in both pLM encoders, reinforcing the importance of window-level embeddings. Moreover, the heatmap from the ProtTrans encoder (see Figure 2a) reveals strong associations with distant residues (token 7, denoted in green dot, shows high associativity with tokens 59 and 63), suggesting its capacity to capture global sequence information more effectively. In contrast, the attention distribution in the CaLM encoder is skewed towards neighboring residues (see Figure 2b), which might explain its relatively poorer performance when used independently. However, when examining the individual heads in the CaLM encoder (see Figure 2c), some heads, such as head 6 and head 8, manage to capture associations with some distant residues, bringing useful information for prediction. Given that the attention heads across each pLM exhibit varied patterns in their distribution of weights, this diversity likely contributes to the enhanced predictive performance observed when the embeddings from both models are combined.

**Figure 2.**
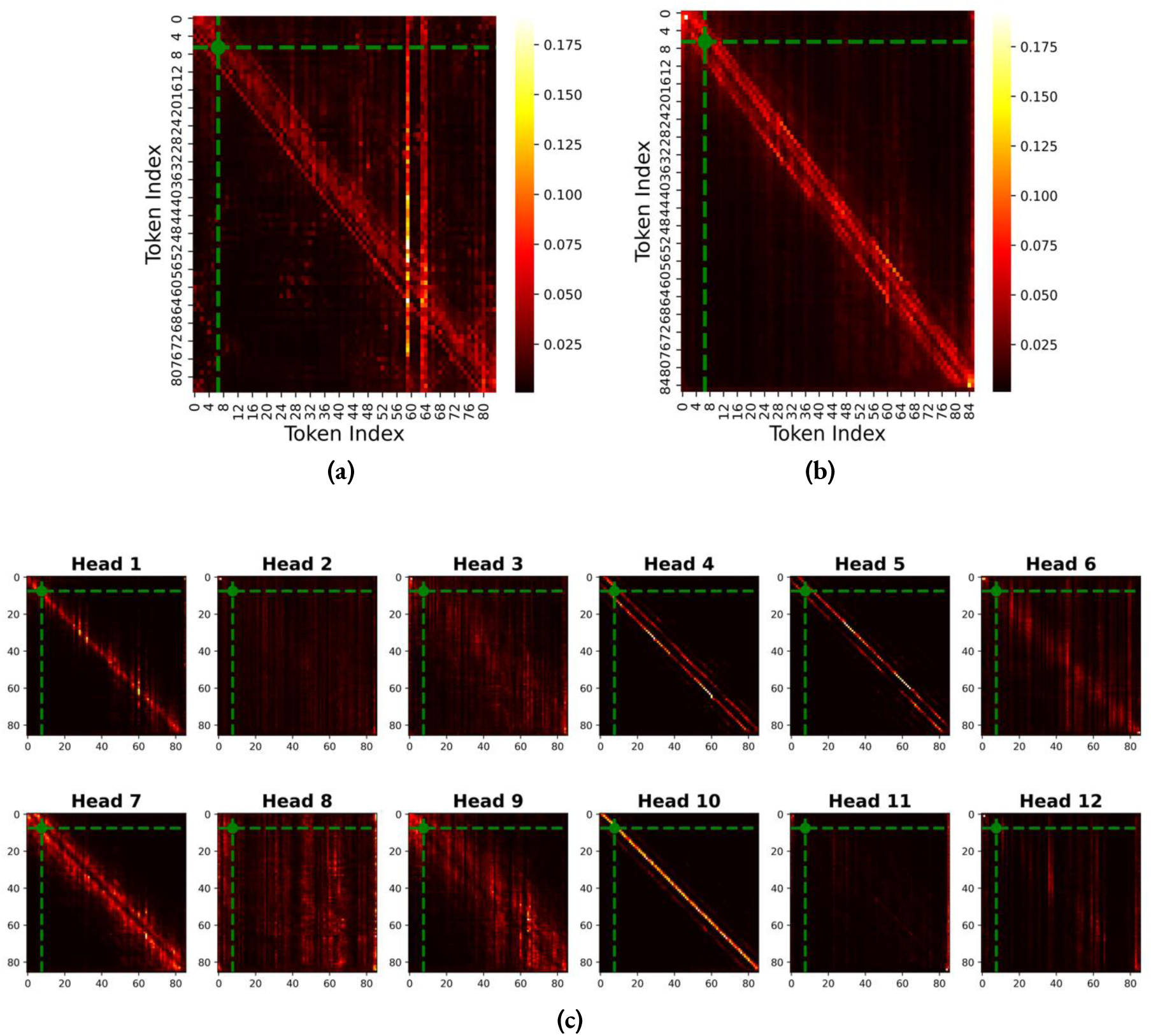
Heatmaps of averaged attention weights over the heads of the last encoder layer of (a) ProtTrans and (b) CaLM with protein P30047 as an input (86 tokens excluding <CLS> and <EOS>). The heatmaps in (c) display the individual attention heads from the last encoder layer of CaLM. The green dot in each denotes an experimentally annotated P-site (at index 7).

**Table 2.** 10-fold cross-validation results (mean ± one standard deviation) for CaLM, ProtTrans, and their fusion (CaLMPhosKAN) for *S+T* and *Y* datasets. The highest values for each metric are bolded in each column.

| Dataset | Embeddings | MCC | PRE | REC | F1<sub>weighted</sub> | AUPR |
| --- | --- | --- | --- | --- | --- | --- |
| Serine +<br>Threonine<br>($S+T$) | CaLM | 0.44±0.01 | 0.74±0.01 | 0.68±0.01 | 0.71±0.01 | 0.80±0.01 |
| | ProtTrans | 0.46±0.01 | 0.75±0.01 | 0.69±0.02 | 0.72±0.01 | 0.81±0.01 |
|  | CaLMPhosKAN | **0.48±0.01** | **0.76±0.01** | **0.70±0.01** | **0.74±0.01** | **0.83±0.01** |
| Tyrosine<br>($Y$) | CaLM | 0.32±0.01 | 0.67±0.01 | 0.60±0.01 | 0.65±0.01 | 0.69±0.01 |
| | ProtTrans | 0.33±0.01 | 0.67±0.01 | 0.62±0.02 | 0.66±0.01 | 0.70±0.01 |
|  | CaLMPhosKAN | **0.34±0.01** | **0.68±0.01** | **0.63±0.02** | **0.67±0.01** | **0.71±0.01** |

### 3.2 Optimal Wavelet Function

Different wavelet functions, each with distinct shapes and mathematical properties, influence how they capture and represent information within the data. In this context, we examined the performance of Wav-KAN models using 10-fold cross-validation across five distinct mother wavelet functions - Morlet, Meyer, Mexican Hat (Ricker), Shannon, and Derivative of Gaussian (DoG). Radar plots (with normalized metrics) for both the *S+T* and *Y* datasets visually illustrate the comparative performance of each wavelet function (see Figure 3). In these plots, the Morlet and Meyer wavelets showed limited coverage, indicating suboptimal performance. Conversely, the Mexican Hat and Shannon wavelets showed greater robustness in performance metrics with considerable coverage. In the *S+T* dataset, Shannon slightly led in AUPR, while in the *Y* dataset, its performance was on par to Mexican Hat (see Supplementary Section S5 for results in tabulated form). Most notably, the DoG wavelet covered the most extensive coverage on the plots (represented in purple), outperforming the others across all evaluated metrics. This observation is consistent with findings from the Wav-KAN paper ^50^, which noted the DoG wavelet’s superior performance over the Mexican Hat and Spline-KAN ^51^ in the MNIST dataset. Consequently, the DoG wavelet was selected for integration into the Wav-KAN module of the CaLMPhosKAN

**Figure 3.**
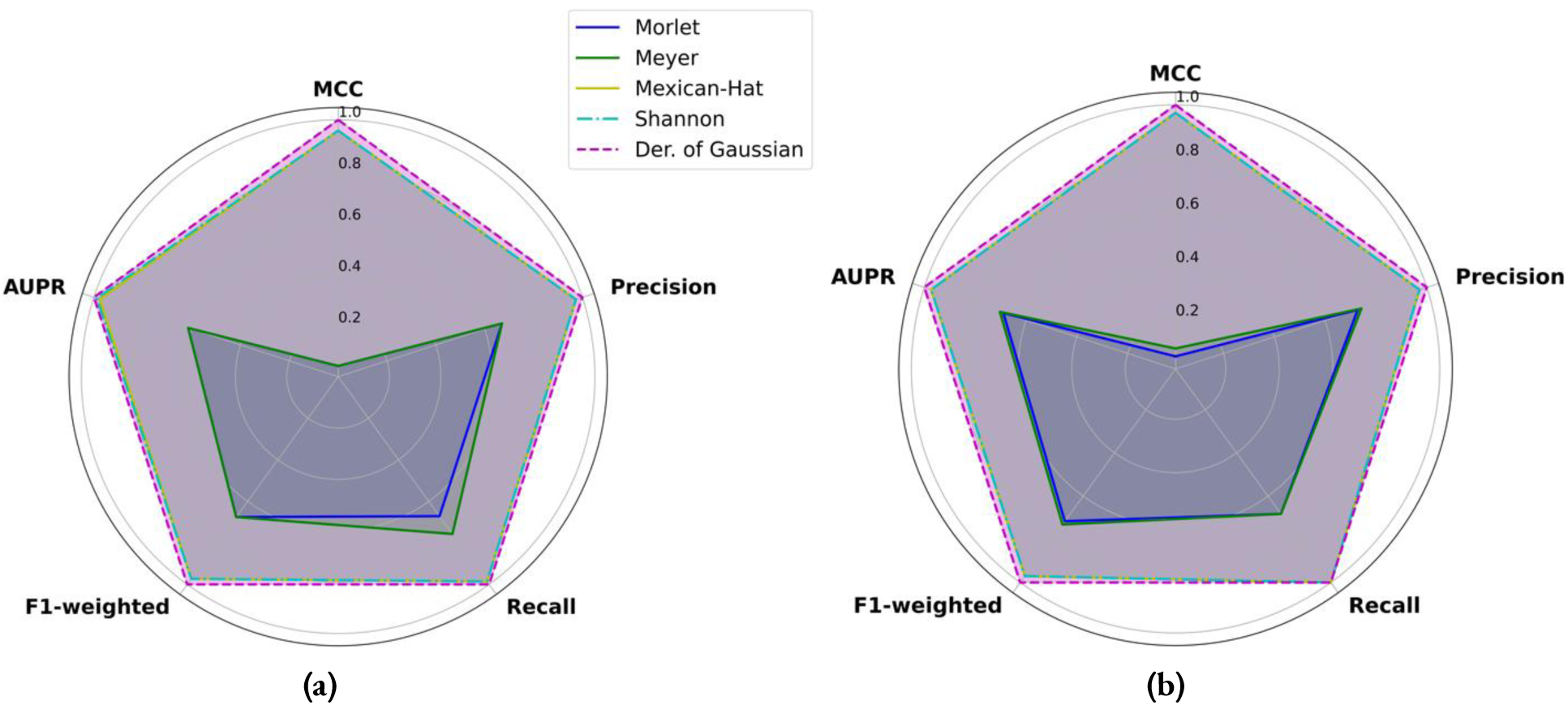
Radar plots comparing five wavelet functions across various performance metrics using 10-fold cross-validation, with each metric normalized between 0 and 1 (using max scaling; each element *x* in a feature column is divided by *max* of that column i.e. *x’= x/ max*) to facilitate direct comparisons across wavelets. Plot (a) on left displays results for the *S+T* dataset and Plot (b) on the right for the *Y* dataset. The wavelets are color-coded as follows: Morlet in deep blue (solid line), Meyer in green (solid line), Mexican Hat in yellow (solid line), Shannon in cyan (dashed line), and Derivative of Gaussian (DoG) in purple (dashed line).

### 3.3 Benchmarking on Independent Test Set

The performance of CaLMPhosKAN was compared on the independent test sets with four existing predictors: DeepPSP ^27^, CapsNet ^17^, Musite ^26^, and MusiteDeep ^26^. For the current best predictor, DeepPSP, we extracted the prediction results that are available on their GitHub repository and computed the performance on samples corresponding to our test set. The performance metrics for the other predictors were adopted from the DeepPSP paper, as our test set is a subset of their test set and the number of sites is similar. This comparison ensures a fair evaluation of CaLMPhosKAN’s performance relative to existing models. It is important to note that the performance on the *S+T* and *Y* datasets were compared separately. Table 3 reports the comparative performance of CaLMPhosKAN along with the four existing predictors based on six measures MCC, PRE, REC, F1, F1_weighted_, and AUPR (see Supplementary Section S6 for PR curves). On the *S+T* dataset, CaLMPhosKAN produced MCC, F1_weighted_, and AUPR of 0.41, 0.51, and 0.53 respectively. CaLMPhosKAN shows improvement on all these three key metrics over the current best predictor DeepPSP. A similar trend was observed in the *Y* dataset, where CaLMPhosKAN registered improvements with MCC, F1_weighted_, and AUPR scores of 0.30, 0.78, and 0.42 respectively. Overall, CaLMPhosKAN showed improvements in all six-performance metrics expect REC.

Additionally, we assess the statistical significance of the difference in the predictive performance of CaLMPhosKAN compared to DeepPSP using McNemar’s chi-squared test. CaLMPhosKAN produced *p*-values of 0.00 (*X*^2^ = 1755.16) and 1.69e^-146^ (*X*^2^ = 664.35) on the *S+T* dataset and *Y* dataset, respectively, which are well below the significance level (***α*** = 0.05). This indicates a statistically significant difference between the performance of CaLMPhosKAN and DeepPSP. All these findings suggest that CaLMPhosKAN is a robust predictor of general phosphorylation sites in proteins.

**Table 3.** Independent test results on *S+T* and *Y* test sets, comparing the performance of CaLMPhosKAN with five other existing predictors. The highest values for each metric are bolded in each column.

| Dataset | Predictors | MCC | PRE | REC | F1 | F1<sub>weighted</sub> | AUPR |
| --- | --- | --- | --- | --- | --- | --- | --- |
| Serine +<br>Threonine<br>($S+T$) | **CaLMPhosKAN** | **0.41** | **0.47** | 0.57 | **0.51** | **0.83** | **0.53** |
|  | DeepPSP | 0.38 | 0.39 | 0.69 | 0.48 | 0.79 | 0.51 |
|  | MusiteDeep | 0.33 | 0.32 | 0.70 | 0.44 | - | 0.46 |
|  | Musite | 0.20 | 0.22 | 0.76 | 0.35 | - | 0.33 |
|  | CapsNet | 0.27 | 0.24 | **0.88** | 0.38 | - | 0.31 |
| Tyrosine<br>($Y$) | **CaLMPhosKAN** | **0.30** | **0.41** | 0.46 | **0.44** | **0.78** | **0.42** |
|  | DeepPSP | 0.26 | 0.32 | 0.66 | 0.42 | 0.70 | 0.39 |
|  | MusiteDeep | 0.20 | 0.35 | 0.35 | 0.35 | - | 0.33 |
|  | Musite | 0.14 | 0.24 | 0.68 | 0.35 | - | 0.28 |
|  | CapsNet | 0.20 | 0.23 | **0.88** | 0.37 | - | 0.19 |

### 3.4 Performance on Intrinsically Disordered Regions

IDRs and non-IDRs exhibit distinct structural and functional properties that can influence phosphorylation. Specifically, IDRs are often enriched in P-sites because their ability to adopt multiple conformations makes them accessible to kinases ^56^. Here, we assess the predictive performance of CaLMPhosKAN specifically in regions of disorder and order. We utilized the popular protein intrinsic disorder region (IDR) prediction tool, flDPnn ^57^, to identify *S* and *T* sites located within disordered and non-disordered regions on proteins in the test set. Subsequently, we segregated the sites based on their regions, disordered and non-disordered, and applied our tool, CaLMPhosKAN, to each set separately. For benchmarking, we compared our results with the current best predictor, DeepPSP, by evaluating its performance on both the disordered and non-disordered datasets. From the bar graphs in Figure 4, it can be observed that CaLMPhosKAN achieves better MCC, F1_weighted_, and AUPR than DeepPSP in both IDR and non-IDR regions. Notably, there appears to be a difference in the performance of these models in inter-regions, which can be attributed to the variation in the distribution of P and NP sites.

**Figure 4.**
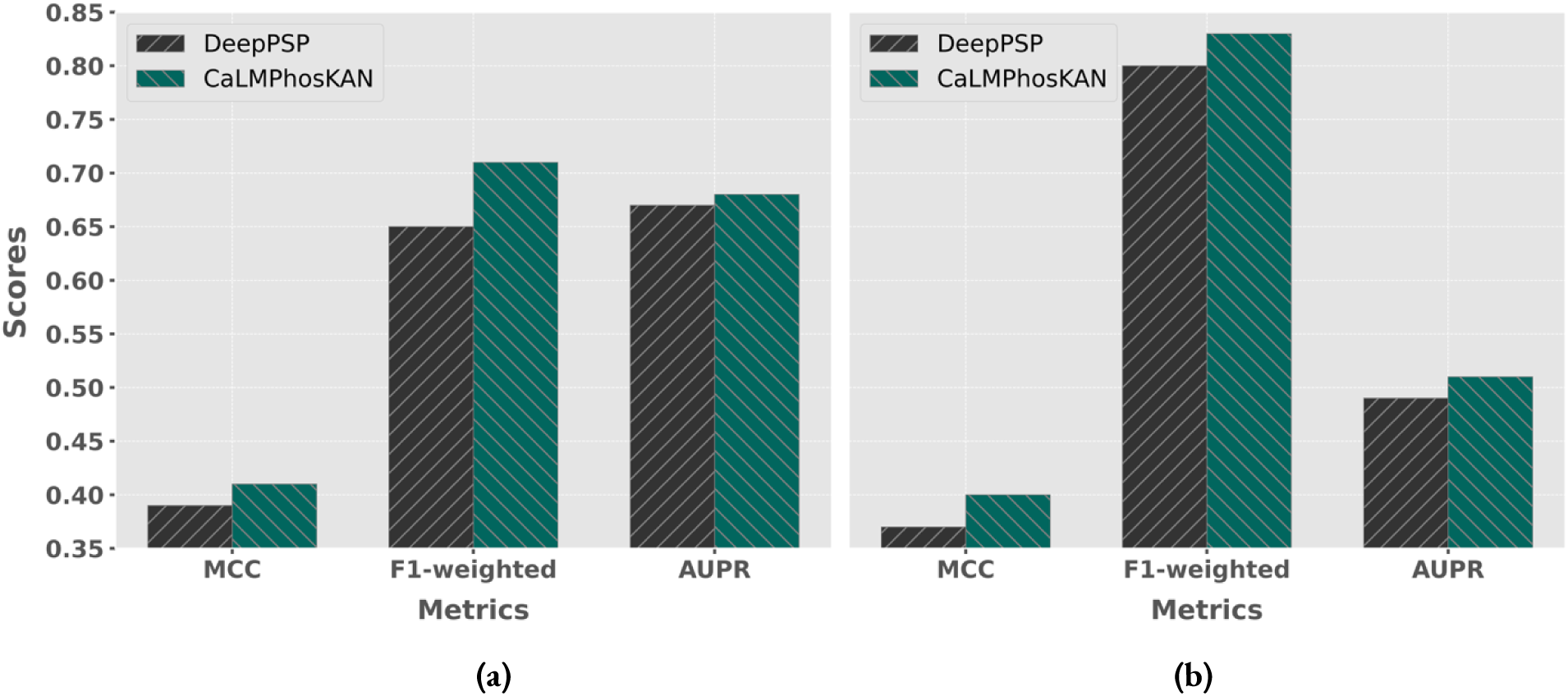
Bar graphs comparing the performance of CaLMPhosKAN and DeepPSP based on MCC, F1, and AUPR for a) IDR regions (left) and b) non-IDR regions (right) of proteins in the test set.

We also plotted the probability distribution of these models on the regions split by ground truth (P vs NP) using violin plots presented in Figure 5. For disordered regions, there is considerable overlap between P and NP sites. The distribution of P sites shows a wide range of scores, indicating variability and some uncertainty in predictions. Similar to DeepPSP, CaLMPhosKAN also shows considerable overlap between NP and P predictions, but the distribution for P sites is somewhat more concentrated than in DeepPSP. For non-disordered regions, in DeepPSP, the NP and P distributions are more distinct than in disordered regions, though there is still overlap. Similarly, in CaLMPhosKAN, the overlap between NP and P predictions is less pronounced compared to disordered regions. Predictions for P-sites are more concentrated towards higher scores, indicating greater confidence in these predictions. Overall, CaLMPhosKAN generally shows more concentrated predictions for P sites in both regions compared to DeepPSP, suggesting higher confidence. Both models, however, present a broader spectrum of predictions for P sites in disordered regions, highlighting the complexity of these areas.

**Figure 5.**
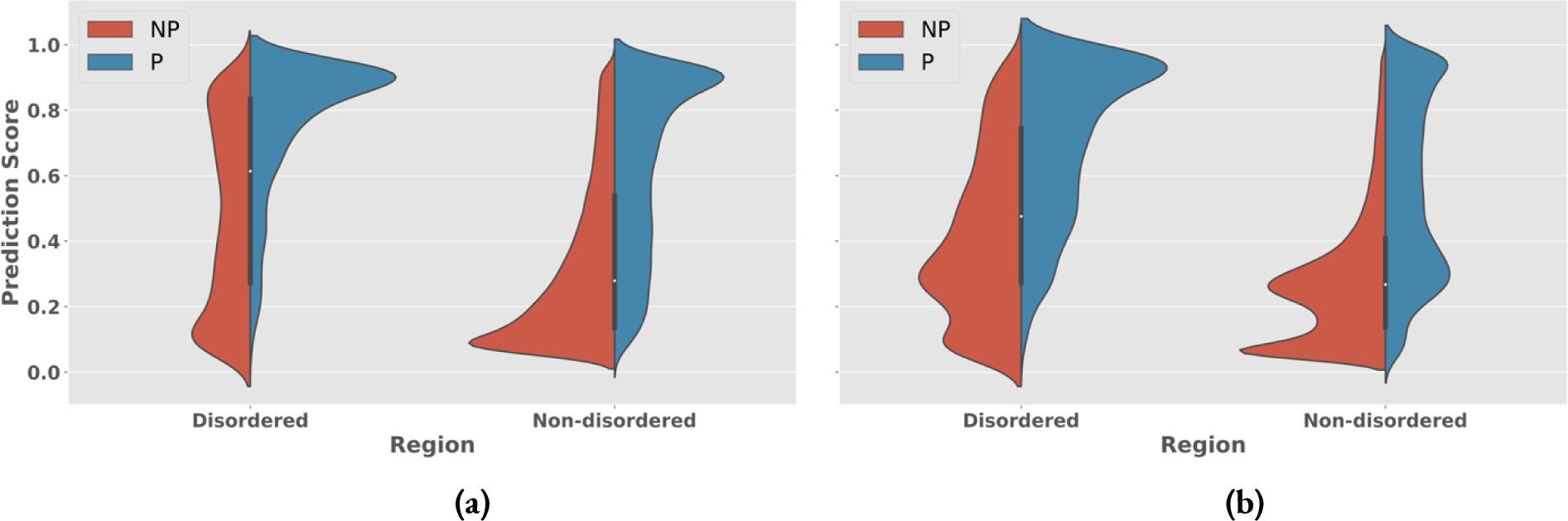
Violin plots showing the distribution of prediction probabilities in disordered and non-disordered regions split by ground truth (P vs NP) for (a) DeepPSP (left) and (b) CaLMPhosKAN (right). The ground truth P is shown in blue while NP in red. The prediction score ranges from 0.0 to 1.0.

Quantitative assessment of the overlap between P and NP predictions using Equation 8 further supports these observations. In non-disordered regions, CaLMPhosKAN shows a slightly better performance with less overlap (1.599) compared to DeepPSP (1.622). Conversely, in disordered regions, the overlap in CaLMPhosKAN is marginally less than in DeepPSP (1.619 vs 1.621). The derivation of Equation 8 is provided below.

Let *P_X,R_* represent the set of predictions for P-sites for model *X* in region *R,* and *N_X, R_* represent the set of predictions for NP-sites for model *X* in region *R*. Let |*P_X, R_*| be the number of positive predictions for model *X* in region *R* and |*N_X, R_*| be the number of negative predictions for model *X* in region *R*.

For each positive prediction *p* ∈ *P_X,R_*, count the number of negative predictions *n* ∈ *N_X, R_* that are less than or equal to *p*. Let *C_P, X, R_* be this count which can be expressed as:

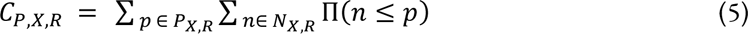

Now, for each negative prediction *n* ∈ *N_X,R_*, count the number of positive predictions *p* ∈ *P_X, R_* that are less than or equal to *n*. Let *C_NP, X, R_* be this count which is expressed as :

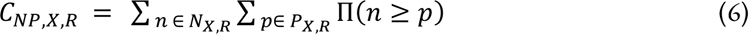

The total overlap *Overlap(X, R)* is the sum of the overlaps from positive and negative predictions normalized by the product of the number of positive and negative predictions:

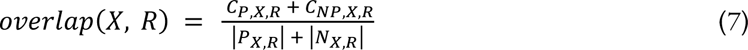

Finally, the general equation of overlap between positive and negative distributions for any model *X* in any region *R* is obtained by combining equations (3), (4), and (5) :

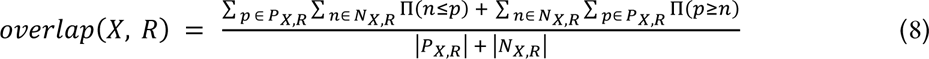

where *Π*(.) is the indicator function, which equals 1 if the condition inside is true, and 0 otherwise.

## 4. Discussion and Conclusion

Phosphorylation is recognized as one of the most pivotal post-translational modifications (PTMs) due to its regulatory roles in myriad cellular processes. The recent surge in utilizing amino acid-aware protein language models (pLMs) has notably advanced bioinformatics tasks. However, the incorporation of codon-level information remains unexplored.

In this work, we introduced a novel approach to phosphosite prediction that incorporates codon-level information by leveraging a protein language model (pLM) trained on codon sequences, named CaLM. We began by translating amino acid sequences into coding sequences through the meticulous mapping of UniProt entries to the NCBI database, employing a dynamic programming-based algorithm to ensure mapping reliability. Codon-aware embeddings were then extracted using the CaLM encoder from these mapped sequences. To enrich the representation, these embeddings were combined with amino acid-aware pLM embeddings through an early fusion strategy. Subsequently, potential phosphorylation sites were characterized at the window level using a local window approach to capture the sequence context. A spatiotemporal model (ConvBiGRU) was then applied to capture important associations between residue embeddings within these window-level representations. The final inference of site of interest was made using a wavelet function-based Kolmogorov–Arnold Network (KAN) learning these associations. The overall framework, termed CaLMPhosKAN, exhibited superior performance in independent testing. Furthermore, cross-validation experiments highlighted the contribution of codon-aware embeddings to the prediction performance. The performance of CaLMPhosKAN was also specifically assessed in intrinsically disordered regions (IDRs) of proteins, providing valuable insights.

CaLMPhosKAN highlights the benefits of integrating codon information into prediction tasks and suggests potential applications in various other bioinformatics inference tasks. Further enhancements to CaLMPhosKAN could include the integration of structural and geometrical representations ^58,59^ (e.g. graph, voxel, etc.) to refine its predictive capabilities. CaLMPhosKAN will soon be made publicly available to the scientific community. This work represents a step forward in the computational prediction of PTM sites by opening new avenues for exploring the functional dynamics of proteins through codon-aware models.

## Key Points

- CaLMPhosKAN utilizes codon-aware embeddings derived from a codon language model, which are fused with amino acid-aware embeddings to enhance feature representation.
- Raw protein sequences are meticulously mapped to their corresponding protein-coding DNA sequences, which are then used as input to the codon language model.
- The sites of interest are represented at the window level, and a ConvBiGRU is employed to extract features that capture spatial and temporal correlations of neighboring residues.
- A wavelet-based Kolmogorov-Arnold Network (KAN) is used as the classification head, leveraging the advantages of wavelet transformations.
- CaLMPhosKAN outperforms other deep learning approaches in independent testing.

## Supporting Information

The supplementary materials will be published soon.

## Author Contributions Statement

P.P. and D.B.K. conceived and designed the experiments. P.P. and C.C. performed all the experiments and data analysis. H.D.I., P.P., and S.P. worked on codon translation. S.P. and M.C. implemented the existing predictors on the independent test sets. S.P., C.C., M.C., and H.D.I. validated all the programs and models. P.P., C.C., S.P., D.B.K., M.C., and H.D.I. wrote and revised the manuscript. D.B.K oversaw the overall project. All authors have read and agreed to the published version of the manuscript.

## Funding

This research was funded by the National Science Foundation (NSF), grant number 1901793, 2210356 (granted to D.B.K.). The high-performance computational resources utilized were made possible by NSF award 2215734 (to D.B.K). Part of this work was supported by the Michigan Department of Health and Human Services (MDHHS) through the Michigan Sequencing and Academic Partnerships for Public Health Innovation and Response (MI-SAPPHIRE) grant.

## Notes

The authors declare no competing financial interest.

## Data and Software availability

The CaLMPhosKAN model will be publicly released soon.

## Acknowledgment

We acknowledge the use of the DeepBlizzard HPC Cluster located at Michigan Tech. We also acknowledge the helpful communication with Dr. Carlos Outeiral (University of Oxford) regarding the codon language model, and with Dr. Lukasz Kurgan (Virginia Commonwealth University) regarding disordered regions.

## Abbreviations

CaLM: Codon Adaptation Language Model
DNA: Deoxyribonucleic Acid
Wav-KAN: Kolmogorov-Arnold Network
ConvBiGRU: Convolutional Gated Recurrent Unit
DoG: Derivative of Gaussian
KAN: Kolmogorov-Arnold Network
mRNA: Messenger RNA
PTM: Post-translational modification
ML: Machine Learning
SENet: Squeeze and excitation Network
CapsNet: Capsule Network
DL: Deep Learning
CDHIT: Cluster Database at High Identity with Tolerance
S: serine
T: threonine
Y: tyrosine
P-sites: phosphorylated/positive sites
NP-sites: non-phosphorylated/negative sites
NCBI: National Center for Biotechnology Information
RefSeq: NCBI Nucleotide Reference sequences
NW: Needleman–Wunsch (NW) algorithm
pLM: Protein Language model
ESM: Evolutionary Sequence Modelling
T5: Text- to-Text Transfer Transformer
MLM: Masked Language Modeling
U: Selenocysteine
Z: Pyrrolysine
O: Hydroxyproline
B: Beta-amino acids
2DCNN: Two Dimensional Convolutional layer
BiGRU: Bidirectional Gated Recurrent Unit
CWT: Continuous Wavelet Transform
DWT: Discrete Wavelet Transform
Dog: Derivative of Gaussian
BCE: Binary-Cross Entropy
MCC: Matthews Correlation Coefficient
PRE: Precision
REC: Recall
AUC: Area Under Curve
AUPR: Area Under the Precision-Recall Curve
AUROC: Area Under Receiver Operating Characteristic
IDRs: Intrinsically Disordered Regions
non-IDRs: Non-Intrinsically Disordered Regions.

## Notes

### Competing Interest Statement

The authors have declared no competing interest.

